# Neutralization against B.1.351 and B.1.617.2 with sera of COVID-19 recovered cases and vaccinees of BBV152

**DOI:** 10.1101/2021.06.05.447177

**Authors:** Pragya D. Yadav, Gajanan N. Sapkal, Raches Ella, Rima R. Sahay, Dimpal A Nyayanit, Deepak Y. Patil, Gururaj Deshpande, Anita M. Shete, Nivedita Gupta, V. Krishna Mohan, Priya Abraham, Samiran Panda, Balram Bhargava

**Affiliations:** Indian Council of Medical Research-National Institute of Virology, Pune, IndiaPin-411021; Bharat Biotech International Limited, Genome Valley, Hyderabad, Telangana, India Pin-500 078; Indian Council of Medical Research, V. Ramalingaswami Bhawan, Ansari Nagar, New Delhi, India Pin-110029

**Keywords:** SARS-CoV-2, Variant of Concern, BBV152 vaccine, B.1.351, B.1.617.2

## Abstract

Recently, multiple SARS-CoV-2 variants have been detected across the globe. The recent emergence of B.1.617 lineage has created serious public health problem in India. The high transmissibility was observed with this lineage which has led to daily increase in the number of SARS-CoV-2 infections. Apparently, the sub-lineage B.1.617.2 has slowly dominated the other variants including B1617.1, B.617.3 and B.1.1.7. With this, World Health Organization has described B.1.617.2 as variant of concern. Besides this, variant of concern B.1.351 has been also reported from India, known to showreducedefficacyfor many approved vaccines. With the increasing threat of the SARS-CoV-2 variants, it is imperative to assess the efficacy of the currently available vaccines against these variants. Here, we have evaluated the neutralization potential of sera collected from COVID-19 recovered cases (n=20) and vaccinees with two doses of BBV152 (n=17) against B.1.351 and B.1.617.2 compared to the prototype B.1 (D614G) variant.The finding of the study demonstrated a reduction in neutralization titers with sera of COVID-19 recovered cases(3.3-fold and 4.6-fold) and BBV152 vaccinees (3. 0 and 2.7 fold) against B.1.351 and B.1.617.2 respectively.Although, there is reduction in neutralization titer, the whole-virion inactivated SARS-CoV-2 vaccine (BBV152) demonstrates protective response against VOC B.1351 and B.1.617.2.

---

Recently, several SARS-CoV-2 variants have emerged from various countries worldwide.^1^ Among them, VOC i.e., B.1.1.7 (Alpha), B.1.351 (Beta), B.1.1.28.1 (Gamma) and B.1.617.2 (Delta) are serious public health threats because of their association with the higher transmissibility and the potential immune escape.^2-4^ Various reports have been published on the neutralization efficacies with the sera of the currently available COVID-19 vaccines against these variants. However, the immune escape of B.1351 variant has been matter of concern for the COVID-19 vaccination program. It has shown resistance to several approved vaccines such as mRNA-1273, BNT162b2, ChAdOx1 nCoV-19, NVX-CoV2373.^5-9^ Another reason of global concern is the recent emergence and detection of highly transmissible B.1.617.2 variant from 44 countries including India.^4^ An inactivated SARS-CoV-2 vaccine, BBV152 is currently rolled out under the national COVID-19 vaccination program in India. The neutralization potential of the BBV152 has been already studied with the B.1, B.1.1.7, B.1.1.28.2 B.1.617.1 and found to be effective against these variants.^10-12^

Here, we assessed the neutralization potential of sera from COVID-19 recovered cases (n=20) post 5-20 weeks of infection and vaccinees 28 days after two doses of BBV152 (n=17) against B.1.351, B.1.617.2 compared to prototype B.1 (D614G). The recovered cases were infected with B.1 (n=17) and B.1.617.1 lineage (n=3). SARS-CoV-2 isolates B.1, B.1.351 and B.1.617.2 were propagated at ICMR-NIV, Pune from the clinical samples using Vero CCL-81 cells and were used for a 50% plaque reduction neutralization test (PRNT50)^13^. The virus genome was confirmed using next generation sequencing for its mutation relevant to assigned lineages as described earlier.^14^ The amino acid changes in the spike region of the B.1 (D614G) (NIV2020-770, GISAID accession number: EPI_ISL_420545), B.1.351 (NIV2021-893, GISAID accession number: EPI_ISL_2036294) and B.1.617.2 (NIV2021-1916, GISAID accession number: EPI_ISL_2400521) are indicated in comparison to Wuhan isolate HU-1(NC_045512.2) in Figure 1. The amino acid changes marked in spike protein for B.1 are D614G; B.1.351 are D80A, D215G, L242del, K417N, E484K, N501Y, D614G, and A701V; and B.1.617.2 are T19R, G142D, E154del, A222V, L452R, T478K, D614G, P681R and D950N.

**Figure 1:**
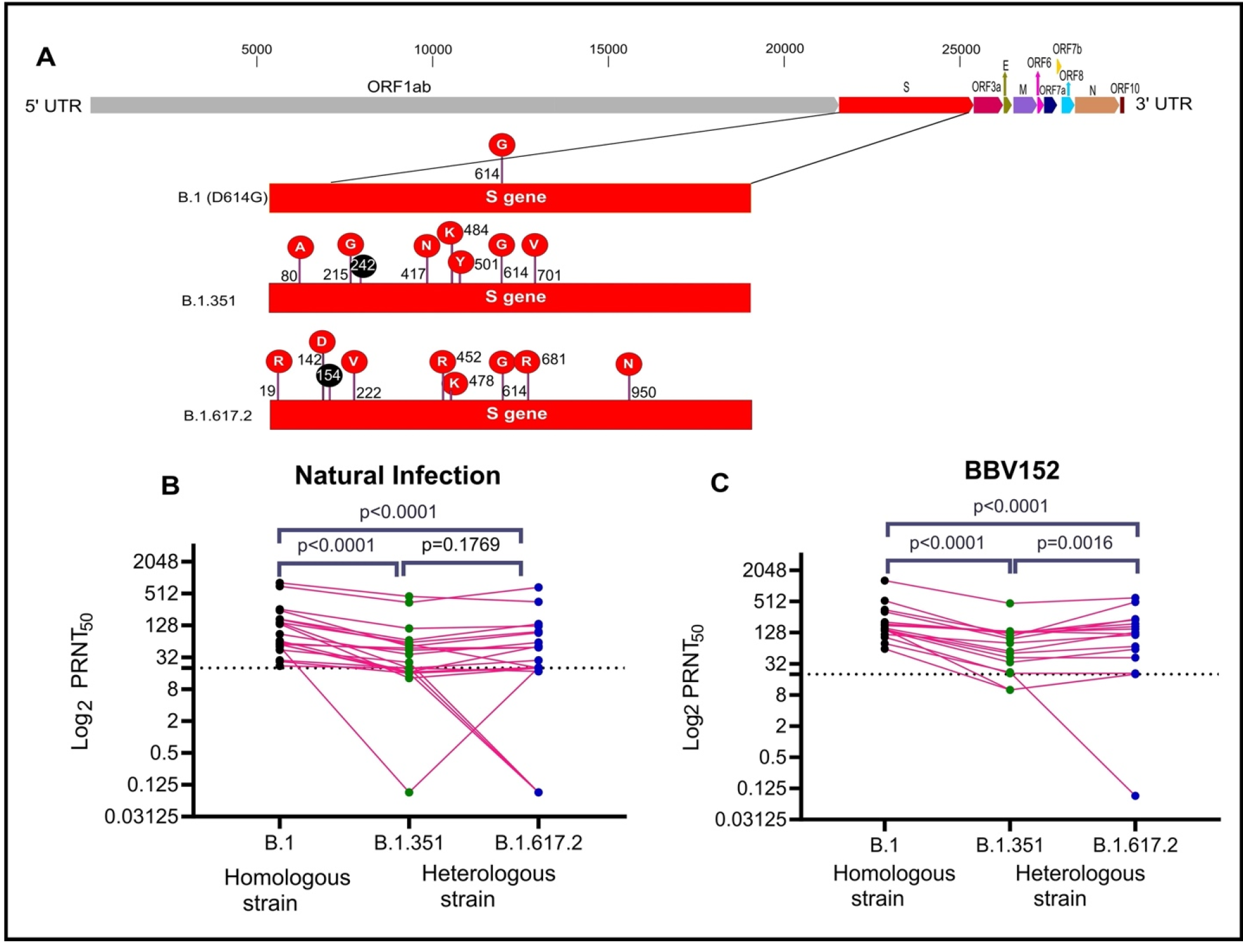
Mutation in Spike protein and neutralization of SARS CoV-2 for B.1.351, B.1.617.2 and B.1: A) The complete genome of SARS-CoV-2 indicting the different genes encoded by it. The inset highlights the amino acid changes in the spike region of the B.1 (D614G) (NIV2020-770, GISAID accession number: EPI_ISL_420545), B.1.351 (NIV2021-893, GISAID accession number: EPI_ISL_2036294) and B.1.617.2 (NIV2021-1916, GISAID accession number: EPI_ISL_2400521) are indicated in comparison to Wuhan isolate HU-1(NC_045512.2). Black color depicts deletion of amino acids. The amino acid changes marked in spike protein for B.1 are D614G; B.1.351 are D80A, D215G, L242del, K417N, E484K, N501Y, D614G, and A701V; and B.1.617.2 are T19R, G142D, E154del, A222V, L452R, T478K, D614G, P681R and D950N. Black color depicts deletion of amino acids. Neutralization titer of naturally infected cases sera (B) and Vaccinees sera (C) against B.1. (Black color), B.1.351 (pink color) and B.1.617.2 (D614G) (Blue color) are shown as matched-pair plot. A paired two-tailed comparison was performed using a two-tailed Wilcoxon matched-pairs signed-rank test with a p-value of 0.05. **** represent p-value <0.0001 and **p value=0.0038, ns= non-significant p-value. The dotted line indicates the detection limit of the test.

A four-fold serial dilution of the serum samples were mixed with an equal amount of each virus suspension (50-60 Plaque forming units in 0.1 ml) separately. After incubating the mixtures at 37°C for 1 hr, each virus-diluted serum sample (0.1 ml) was inoculated onto duplicate wells of a 24-well tissue culture plate containing a confluent monolayer of Vero CCL-81 cells. After incubating the plate at 37°C for 60 min, an overlay medium consisting of 2% Carboxymethyl cellulose (CMC) with 2% fetal calf serum (FCS) in 2× MEM was added to the cell monolayer, and the plate was further incubated at 37°C in 5% CO 2 for 5 days. Plates were stained with 1% amido black for an hour. Antibody titers were defined as the highest serum dilution that resulted in >50 (PRNT50) reduction in the number of plaques.^15^

Geometric mean titer (GMT) for vaccinees sera against B.1, B.1.351 and B.1.617.2 were found to be 187.5 (95%CI: 129.3-271.9), 61.57 (95%CI: 36.34-104.3) and 68.97(95%CI: 24.72-192.4) respectively. The GMT ratio of B.1 to B.1.351 and B.1.617.2 was 3.0 (95%CI: 2.6-3.6) and 2.7 (95% CI: 1.4-5.2).

Similarly, GMT titers in sera of recovered cases against B.1, B.1.351, and B.1.617.2 were 97.8 (95%CI: 61.2-156.2), 29.6 (95%CI: 13.4-65.0) and 21.2(95% CI: 6.4-70.1) respectively. The GMT ratio of B.1 to B.1.351 and B.1.617.2 was 3.3 (95%CI: 2.4-4.5) and 4.6 (95% CI: 2.2-9.5). Sera of vaccinees and recovered cases had shown a significant reduction in neutralization titer for B.1.351 and B.1.617.2 in comparison to B.1 (p-value :< 0.0001) (Figure 1).

We observed a reduction in neutralization titers with sera of COVID-19 recovered cases and BBV152 vaccines against B.1.351 and B.1.617.2 respectively. Several studies have demonstrated the reduction in the neutralization efficacy with the sera of naturally infected cases and individuals vaccinated with BBIBP-CorV (1.6×), BNT162b2 (6.5×), mRNA-1273 (8.6×), ChAdOx1 nCoV-19 (86×) against B.1351.^1,16,17^ Reduced neutralization with the vaccinees sera of BNT162b2 mRNA (7×) and one dose of ChAdOx1 nCoV-19 was observed against B.1.617.2.^4^

Our study demonstrated that despite a reduction in neutralization titers with BBV152 vaccinees sera against B.1.351 and B.1.617.2, its neutralization potential is well established. Lastly, the broad epitope coverage of an inactivated vaccine (BBV152) decreases the magnitude of reduced neutralization against emerging variants.

## Ethical approval

The study was approved by the Institutional Biosafety Committee and Institutional Human Ethics Committee of ICMR-NIV, Pune, India under the project ‘Propagation of new SARS-CoV-2 variant isolate and characterization in cell culture and animal model’.

## Author Contributions

PDY, GS and PA contributed to study design, data collection, data analysis, interpretation and writing and critical review. RRS, DAN, DYP, GD and AS contributed to data collection, interpretation, writing and critical review. NG, SP, VKM and BB contributed to critical review and finalization of the paper.

## Conflicts of Interest

Authors do not have a conflict of interest among themselves.

## Financial support & sponsorship

Financial support was provided by the Indian Council of Medical Research (ICMR), New Delhi at ICMR-National Institute of Virology, Pune under intramural funding ‘COVID-19’.

## Acknowledgement

Authors gratefully acknowledge the staff of ICMR-NIV, Pune including Mr. Prasad Sarkale, Mr. ShreekantBaradkar, Mr. Hitesh Dighe, Mrs. Savita Patil, Mrs. Tripanra Majumdar, Ms. AashaSalunkhe and Mr. Chetan Patil for extending excellent technical support.

## Notes

### Competing Interest Statement

The authors have declared no competing interest.

## References

1. Abdool Karim SS, de Oliveira T. New SARS-CoV-2 variants—clinical, public health, and vaccine implications. N Engl J Med. 2021.

2. Preliminary genomic characterisation of an emergent SARS-CoV-2 lineage in the UK defined by a novel set of spike mutations. https://virological.org/t/preliminary-genomic-characterisation-of-an-emergent-sars-cov-2-lineage-in-the-uk-defined-by-a-novel-set-of-spike-mutations/563

3. Tegally H, Wilkinson E, Giovanetti M, et al. Emergence and rapid spread of a new severe acute respiratory syndrome-related coronavirus 2 (SARS267 CoV-2) lineage with multiple spike mutations in South Africa. medRxiv. 2020.

4. Planas D, Veyer D, Baidaliuk A, et al. Reduced sensitivity of infectious SARS-CoV-2 variant B. 1.617.2 to monoclonal antibodies and sera from convalescent and vaccinated individuals. bioRxiv. 2021.

5. Madhi SA, Baillie V, Cutland CL, et al. Efficacy of the ChAdOx1 nCoV-19 Covid-19 Vaccine against the B.1.351 Variant. N Engl J Med. 2021

6. Shen X, Tang H, Pajon R, et al. Neutralization of SARS-CoV-2 Variants B. 1.429 and B. 1.351. N Engl J Med. 2021.

7. Xie X, Liu Y, Liu J, et al. Neutralization of SARS-CoV-2 spike 69/70 deletion, E484K and N501Y variants by BNT162b2 vaccine-elicited sera. Nature Med. 2021; 27(4):620–1.

8. Hoffmann M, Arora P, Groß R, et al. SARS-CoV-2 variants B. 1.351 and P. 1 escape from neutralizing antibodies. Cell. 2021.

9. Wang GL, Wang ZY, Duan LJ, et al. Susceptibility of Circulating SARS-CoV-2 Variants to Neutralization. N Engl J Med. 2021.

10. Sapkal GN, Yadav P, Ella R, et al. Inactivated COVID-19 vaccine BBV152/COVAXIN effectively neutralizes recently emerged B.1.1.7 variant of SARS-CoV-2, J Travel Med. 2021; 28 (4): taab051.

11. Sapkal G, Yadav PD, Ella R, et al. Neutralization of B. 1.1. 28 P2 variant with sera of natural SARS-CoV-2 infection and recipients of inactivated COVID-19 vaccine Covaxin. J Travel Med. 2021; taab077.

12. Yadav P, Sapkal GN, Abraham P, et al. Neutralization of variant under investigation B. 1.617 with sera of BBV152 vaccinees. Clin Infect Dis. 2021; ciab411. doi: 10.1093/cid/ciab411.

13. Yadav P, Sarkale P, Razdan A, et al. Isolation and characterization of SARS-CoV-2 VOC, 20H/501Y. V2, from UAE travelers. bioRxiv. 2021.

14. Yadav PD, Nyayanit DA, Shete AM, et al. Complete Genome Sequencing of Kaisodi Virus Isolated from Ticks in India Belonging to Phlebovirus Genus, Family Phenuiviridae. Ticks Tick Borne Dis. 2019, 10 (1), 23–33.

15. Deshpande GR, Sapkal GN, Tilekar BN, et al. Neutralizing antibody responses to SARS-CoV-2 in COVID-19 patients. Ind J Med Res. 2020; 152(1-2):82.

16. Wibmer CK, Ayres F, Hermanus T, et al. SARS-CoV-2 501Y. V2 escapes neutralization by South African COVID-19 donor plasma. Nature Med. 2021; 27(4):622–5.

17. Wu K, Werner AP, Koch M, et al. Serum neutralizing activity elicited by mRNA-1273 vaccine. N Engl J Med. 2021; 384(15):1468–70.

